# Amphetamine Maintenance Therapy During Intermittent Cocaine Self-Administration in Rats: Reduction of Addiction-like Behavior is Associated with Attenuation of Psychomotor and Dopamine Sensitization

**DOI:** 10.1101/2020.01.10.900852

**Authors:** Florence Allain, Benoît Delignat-Lavaud, Marie-Pierre Beaudoin, Vincent Jacquemet, Terry E. Robinson, Louis-Eric Trudeau, Anne-Noël Samaha

## Abstract

**Background:** D-amphetamine maintenance therapy shows promise as a treatment for people with cocaine addiction. Preclinical studies using Long Access (LgA) cocaine self-administration procedures suggest D-amphetamine may act by preventing tolerance to cocaine’s effects at the dopamine transporter (DAT). However, Intermittent Access (IntA) cocaine self-administration better reflects human patterns of use, is especially effective in promoting addiction-relevant behaviors, and instead of tolerance, produces psychomotor, incentive, and neural sensitization. We asked, therefore, how D-amphetamine maintenance during IntA influences cocaine use and cocaine’s potency at the DAT.

**Methods:** Male rats self-administered cocaine intermittently (5 minutes ON, 25 minutes OFF x 10) for 14 sessions, with or without concomitant D-amphetamine (5 mg/kg/day via s.c. osmotic minipump). In Experiment 1, psychomotor sensitization, responding for cocaine under a progressive ratio schedule, responding under extinction and cocaine-primed relapse were assessed. In Experiment 2, rats self-administered cocaine or saline intermittently, with or without D-amphetamine, and the ability of cocaine to inhibit dopamine uptake in the nucleus accumbens core was assessed using fast scan cyclic voltammetry *ex vivo*.

**Results:** IntA cocaine self-administration produced psychomotor sensitization, strong motivation to take and seek cocaine, and it increased cocaine’s potency at the DAT. The co-administration of D-amphetamine suppressed both the psychomotor sensitization and high motivation for cocaine produced by IntA experience, and also reversed sensitization of cocaine’s actions at the DAT, leaving baseline DAT function unchanged.

**Conclusions:** Treatment with D-amphetamine might reduce cocaine use by preventing sensitization-related changes in cocaine potency at the DAT, consistent with an incentive-sensitization view of addiction.

## INTRODUCTION

Several pharmacological approaches are presently under study for treating cocaine addiction, but none is approved by North American, European or other agencies (1–3). One promising strategy is to substitute cocaine with another dopaminergic agent (4), such as D-amphetamine, but in a slower and longer-acting formulation that would be potentially less harmful. Low-dose D-amphetamine effectively decreases cocaine use in humans (5–8), non-human primates (9–13) and rats (14–17), with no or only transient effects on responding for food (9, 10, 12–14). However, it is not entirely clear how D-amphetamine produces these effects, but they do not involve reduced brain cocaine concentrations, cross-tolerance or increases in cocaine’s anxiogenic, or other toxic effects (10, 14). Cocaine blocks dopamine uptake at the dopamine transporter (DAT) to enhance dopamine transmission and produce reward (18), and therefore, D-amphetamine might interfere with cocaine’s actions at the DAT, thereby attenuating addiction-relevant effects (19–22).

Indeed, Siciliano et al. (2018) reported that, in rats, continuous, low-dose D-amphetamine (5 mg/kg/day for 14 d via s.c. minipump) reduces cocaine self-administration by preventing molecular changes that lead to a decrease (tolerance) in the ability of cocaine to inhibit dopamine uptake at the DAT in the nucleus accumbens core (NAcC) (23). However, Long-Access (LgA) cocaine self-administration procedures were used, which produce high and sustained brain cocaine concentrations (24–26). In humans, cocaine use is typically more intermittent, which would produce peaks and troughs in brain cocaine concentrations (19, 27). In rats, Intermittent Access (IntA) cocaine self-administration results in much less cumulative cocaine intake than LgA, but in many respects is more effective in producing addiction-like behavior than LgA [(24–26, 28–33); Reviewed in (19, 20)]. Of particular relevance here, LgA and IntA cocaine experience also produce opposite effects on the dopamine system: tolerance vs. sensitization, respectively, to cocaine-induced inhibition of dopamine uptake at the DAT and cocaine-induced dopamine overflow in the NAcC (33, 34). Our objective here, therefore, was to assess the effect of D-amphetamine treatment during IntA cocaine self-administration on both addiction-relevant behaviors and cocaine’s potency at the DAT.

## METHODS AND MATERIALS

See Supplement for information on subjects, surgeries, self-administration training and statistical analyses.

### Experiment 1: Effects of D-amphetamine Maintenance during Intermittent Cocaine Self-administration on the Development of Psychomotor Sensitization and on Drug-Taking and -Seeking Behaviors

#### Intermittent-Access (IntA) sessions

**Figure 1** (**Exp-1**) shows the sequence of experimental events. After self-administration training (1-h sessions), male Wistar rats were allocated to a ‘COC + A’ group and a ‘COC’ group. A minipump filled with D-amphetamine (5 mg/kg/day) was implanted subcutaneously in the COC + A rats (n = 11). COC rats received a saline-filled minipump (n = 11), or a sham surgery (n = 11). All rats then self-administered cocaine (0.25 mg/kg/injection, over 5 s) under a fixed ratio 3 schedule, for 14 IntA-sessions given on consecutive days. Each 5-h session consisted of ten, 5-min cocaine-available periods separated by 25-min, no cocaine-available periods during which the levers were retracted (24). We measured cocaine intake and locomotion during IntA-sessions.

**Figure 1.**
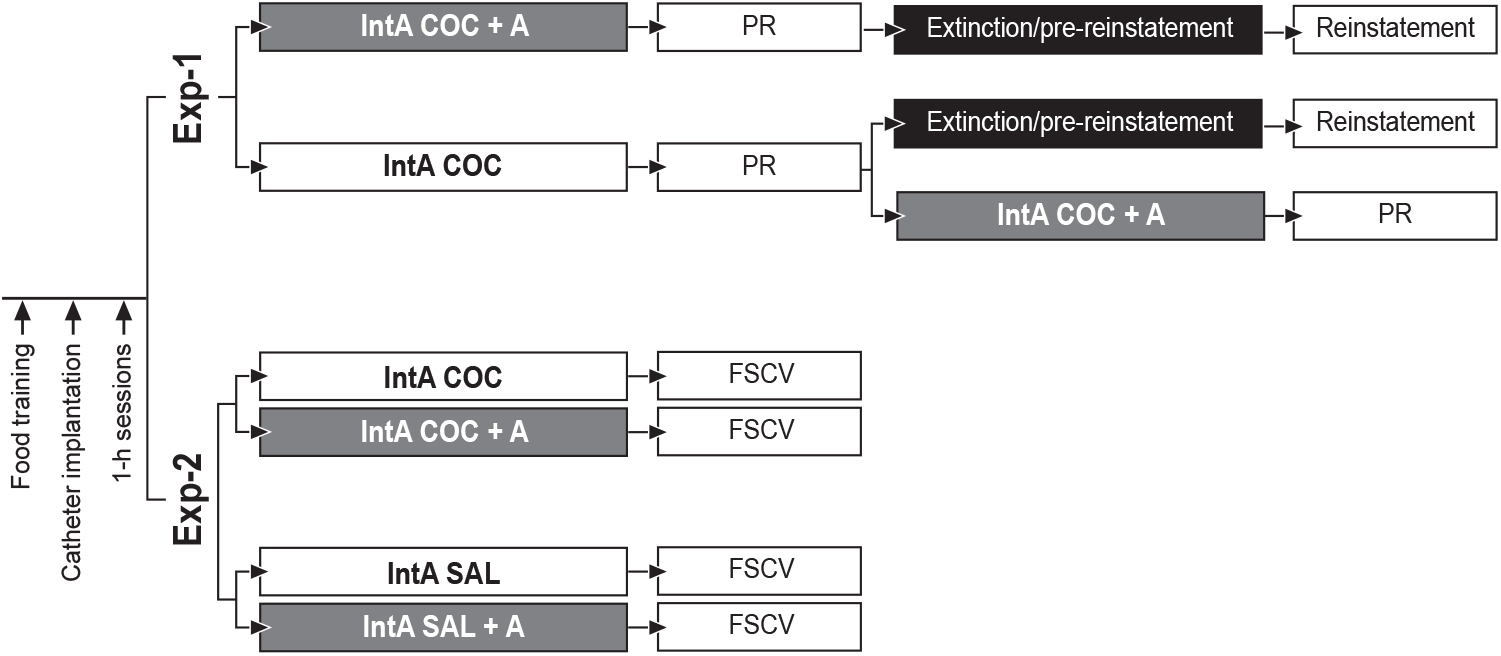
The sequence of experimental events. In experiment 1 (Exp-1), male Wistar rats self-administered cocaine under intermittent-access (IntA) conditions (COC). Some rats received D-amphetamine maintenance (COC + A; 5 mg/kg/day, via s.c. osmotic minipump) during intermittent cocaine self-administration. We assessed cocaine intake, changes in locomotor activity during self-administration sessions, incentive motivation for cocaine [as measured by responding for the drug under a progressive ratio (PR) schedule of reinforcement], and cocaine-induced reinstatement of extinguished cocaine-seeking behavior. A subset of COC rats was also used to compare responding for cocaine under a PR schedule before and after D-amphetamine treatment, in a within-subjects design. In experiment 2 (Exp-2), male Wistar rats self-administered cocaine (COC) or saline (SAL), with or without concomitant D-amphetamine treatment (COC + A and SAL + A). Using fast-scan cyclic voltammetry (FSCV), we then assessed cocaine-induced dopamine reuptake inhibition at the dopamine transporter in the nucleus accumbens core. IntA, Intermittent Access. COC, Cocaine. A, D-amphetamine.

#### Responding under a progressive ratio (PR) schedule of cocaine reinforcement

The day after the last IntA-session, minipumps were removed, and sham rats in the COC group that did not have minipumps received a second sham surgery. Two days later, incentive motivation for cocaine was assessed by quantifying responding for cocaine under a PR schedule [(35), also see Supplement].

#### Extinction

After testing under PR, all COC + A and half of the COC rats were given 10, 2-h extinction sessions (1 session/day), and then five, 2-h pre-reinstatement sessions to further decrease the influence of cocaine cues on subsequent reinstatement testing (36–38). During pre-reinstatement sessions, lever pressing no longer produced any exteroceptive cocaine-associated cues (also see Supplement).

#### Cocaine-induced reinstatement

After extinction and pre-reinstatement sessions, the rats were tested for cocaine-induced reinstatement of drug-seeking behavior. This occurred ~3 weeks after the discontinuation of D-amphetamine treatment. Sessions were similar to pre-reinstatement sessions, except that immediately before each session, rats received cocaine i.p. (0, 7.5 or 15 mg/ml/kg, doses in ascending order, 1 dose/session, within-subjects design). Cocaine seeking was measured as the number of presses on the previously cocaine-associated lever (39, 40). To confirm that lever-pressing behavior returned to baseline between tests, rats received an extinction session with no i.p. injection between reinstatement sessions.

#### Effects of D-amphetamine maintenance after a history of intermittent cocaine intake

We also compared the response to cocaine before and after D-amphetamine treatment, using a within-subjects design. We used half of the COC rats for this (11/22 COC rats; PR scores were similar in the two subsets of COC rats, see **Figure S1**). As **Figure 1** (**Exp-1**) shows, after 14 IntA-sessions and testing under a PR schedule, D-amphetamine-containing minipumps (5 mg/kg/day) were implanted in 11 COC rats, and rats were given 14 additional IntA-sessions, now with concomitant D-amphetamine. The day after the last IntA-session, minipumps were removed. Two days later, the rats were tested under a PR schedule once again, as described above.

### Experiment 2: Effects of D-amphetamine Maintenance during Intermittent Cocaine Self-administration on Cocaine’s Potency at the Dopamine Transporter

A new cohort of rats with D-amphetamine-containing minipumps (5 mg/kg/day) or sham-operated self-administered cocaine (n = 13, including 7 D-amphetamine-treated rats and 5 D-amphetamine-naive rats) or saline (n = 10, including 5 D-amphetamine-treated rats) during 14 IntA-sessions (**Figure 1**; **Exp-2**). This yielded 4 groups; ‘COC’, ‘COC + A’, ‘SAL’, and ‘SAL + A’. The day after the last session, minipumps were removed (sham rats were sham operated). Five days later, brain sections were prepared for FSCV, to measure cocaine-induced inhibition of dopamine uptake in the NAcC (see Supplement).

## RESULTS

### Experiment 1: Effects of D-amphetamine Maintenance during Intermittent Cocaine Self-administration on the Development of Psychomotor Sensitization and on Drug-Taking and -Seeking Behaviors

#### Intermittent Access sessions

Sham rats and rats with a saline minipump were similar on all behavioral measures, and therefore were pooled to form one group (‘COC’). COC and COC + A rats self-administered a similar number of cocaine injections, and both groups also escalated their intake over time (Session effect, F_13_, _403_ = 6.43, *p* < 0.0001; **Figure 2A**). This confirms reports that IntA cocaine experience promotes escalation of intake [(26, 28, 31, 32, 41), but see (24, 25)].

**Figure 2.**
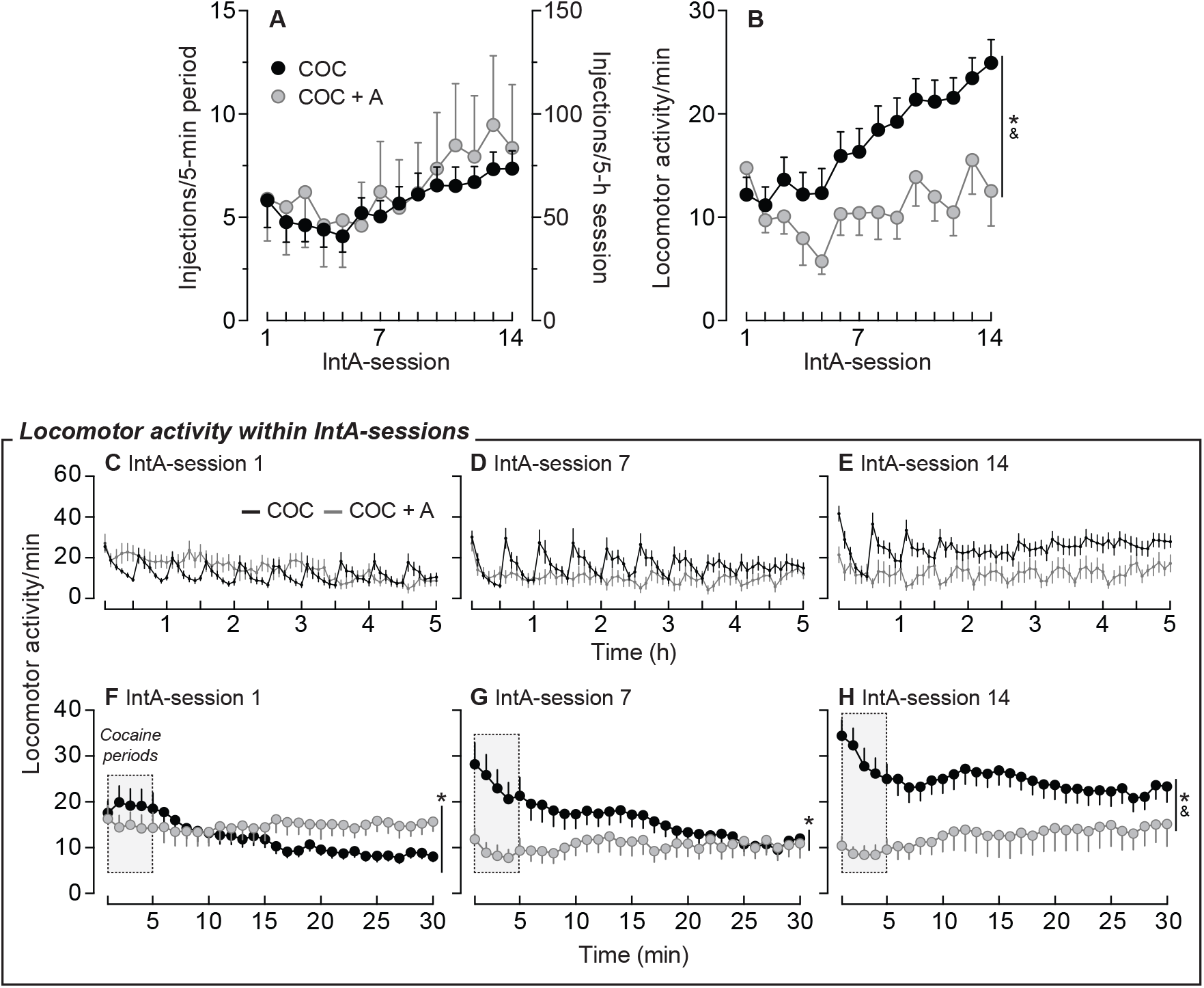
D-amphetamine did not significantly influence cocaine intake, but it abolished both cocaine-induced psychomotor sensitization and spikes in locomotor activity during cocaine self-administration sessions. (**A**) Average number of cocaine injections per 5-min cocaine period (left Y-axis) and total number of cocaine injections per 5-h IntA-session (right Y-axis). (**B**) Locomotor activity/min increased over IntA-sessions only in COC rats, suggesting that only COC rats developed psychomotor sensitization to self-administered cocaine. (**C-E**) Intermittent cocaine intake produced spikes in locomotor activity during the 5-h IntA-sessions, and D-amphetamine suppressed this effect. (**F-H**) Locomotor activity/min averaged over the ten, 5-min cocaine (shaded in gray) and the ten, 25-min no cocaine periods of the 1st, 7th and the 14th IntA-sessions. Data are mean ± SEM. n = 22 for the COC group, and n = 11 for the COC + A group. *P’s < 0.05, Group x Session or Group x Time interaction effect. ^&^P’s < 0.05, Group effect. IntA, Intermittent Access. COC, Cocaine. A, D-amphetamine.

During the 1^st^ IntA-session, locomotion was similar in COC and COC + A rats, but it increased over time only in COC rats, and by the 14^th^ IntA-session, locomotion was greater in COC rats (Group x Session interaction effect, F_13_, _403_ = 5.19, *p* < 0.0001; Group effect, F_1_, _31_ = 4.51, *p* = 0.04; Bonferonni’s tests, *p* = 0.005 at the 14^th^ IntA-session; **Figure 2B**). Increased locomotion over sessions in COC rats could involve increased cocaine intake over sessions. This is unlikely because neither the degree of escalation nor cumulative cocaine intake predicted the increase in locomotion over sessions (*r*^*2*^ = 0.01 and *r*^*2*^ = 0.002, respectively; All *P’*s > 0.05; data not shown). Thus, psychomotor sensitization developed to self-administered cocaine [see also (25, 28, 42)], and D-amphetamine prevented this sensitization.

D-amphetamine also produced qualitative changes in cocaine-induced locomotion. In COC rats, locomotion during IntA-sessions showed a spiking pattern (**Figures 2C-E**), and D-amphetamine attenuated this (**Figures 2C-E**). This is highlighted in **Figures 2F-H**, where locomotion was averaged over the ten, 30-min cycles of cocaine-available (5 min; highlighted with gray shading) and no cocaine-available (25 min) periods. In COC rats only, locomotion increased over sessions (Group x Session effect, F_2, 62_ = 13.96, *p* < 0.0001; **Figures 2F-H**), and was highest during the 5-min cocaine periods (Group x Time effect, F_29, 899_ = 4.45, *p* < 0.0001; **Figures 2F-H**). Locomotion was also greater in COC vs COC + A rats (Group x Time interaction effect; F_29, 899_ = 4.92, *p* < 0.0001, **Figure 2F**; F_29, 899_ = 3.14, *p* < 0.0001, **Figure 2G**; F_29, 899_ = 1.54, *p* = 0.04, **Figure 2H**. Group effect; F_1, 31_ = 9.79, *p* = 0.004; **Figure 2H**; No other comparisons were significant). Thus, D-amphetamine prevented psychomotor sensitization to cocaine and changed the temporal kinetics of cocaine-induced locomotion.

#### Responding under a PR schedule

D-amphetamine treatment was discontinued after the 14^th^ IntA-session. Two days later, rats received PR tests (**Figure 3**). Both groups responded more for higher cocaine doses (Dose effect, F_2, 62_ = 34.11, *p* < 0.0001; **Figure 3A**), and COC rats earned more cocaine than COC + A rats (Group effect, F_1, 31_ = 11.24, *p* = 0.002; **Figure 3A**). This is further illustrated in **Figures 3B-D**, showing cumulative responding for cocaine during PR sessions. COC rats also persisted in responding for cocaine for longer than COC + A rats (Group effect on session duration, F_1, 31_ = 8.51, *p* = 0.007; data not shown). Thus, under a PR schedule, COC rats persevered in working for cocaine as cost in effort increased, to a greater extent than COC + A rats. This suggests that prior D-amphetamine maintenance reduced subsequent incentive motivation to obtain cocaine, perhaps by attenuating the incentive sensitization produced by IntA experience [present study and (24, 26, 30–32)].

**Figure 3.**
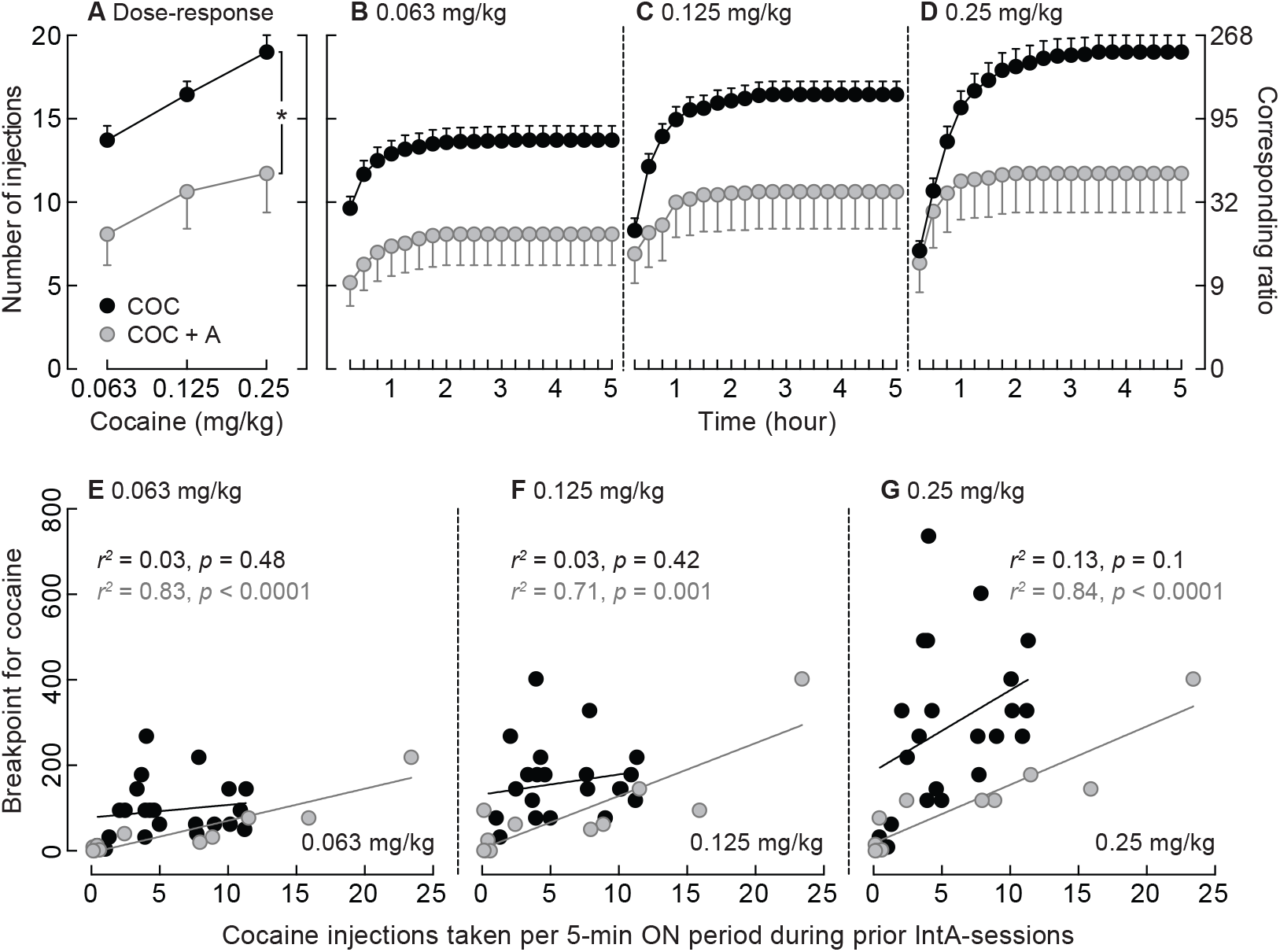
D-amphetamine maintenance during intermittent cocaine self-administration decreased subsequent responding for cocaine under a progressive ratio schedule of reinforcement. (**A**) Across cocaine doses, COC + A rats responded less for cocaine compared to COC rats. (**B-D**) Cumulative number of self-administered cocaine injections (left Y-axis) and corresponding ratio (right Y-axis) during each 5-h progressive ratio test, as a function of cocaine dose. (**E-G**) Correlations between the average number of cocaine injections taken per 5-min cocaine period over the 14 IntA-sessions and the ratio reached during progressive ratio tests at 0.063 mg/kg/infusion (**E**), 0.125 mg/kg/infusion (**F**) and 0.25 mg/kg/infusion (**G**) cocaine. Cocaine intake during IntA-sessions predicted responding for cocaine under a progressive ratio schedule in COC + A rats, but not in COC rats. *p = 0.002, Group effect. Data are mean ± SEM. n = 22 for the COC group, and n = 11 for the COC + A group. IntA, Intermittent Access. COC, Cocaine. A, D-amphetamine.

In COC rats, cocaine intake during IntA-sessions did not predict responding for the drug under a PR schedule [0.063 mg/kg, *r*^*2*^ = 0.03; **Figure 3E**; 0.125 mg/kg, *r*^*2*^ = 0.03; **Figure 3F**; 0.25 mg/kg, *r*^*2*^ = 0.13; **Figure 3G**; All *P*’s > 0.05; See also (25, 26, 43)]. This extends the idea that cocaine consumption and appetitive responding for cocaine are dissociable (24, 26, 43–45). However, in COC + A rats, less cocaine intake during IntA-sessions predicted lower responding under a PR schedule (0.063 mg/kg, *r*^*2*^ = 0.83; **Figure 3E**; 0.125 mg/kg, *r*^*2*^ = 0.71; **Figure 3F**; 0.25 mg/kg, *r*^*2*^ = 0.84; **Figure 3G**; All *P*’s ≤ 0.001). Thus, while D-amphetamine did not reduce cocaine intake during IntA-sessions (**Figure 2A**), the amount of cocaine taken while D-amphetamine is onboard predicts later incentive motivation for cocaine.

#### Extinction

Both groups decreased responding over extinction sessions (Session effect, F_9_, _171_ = 15.37, *p* < 0.0001; **Figure 4A**), but COC rats responded more than COC + A rats (Session x Group effect, F_9_, _171_ = 3.87, *p* = 0.0002; Group effect, F_1_, _19_ = 10.41, *p* = 0.004; **Figure 4A**), especially on the 1^st^ session (Bonferroni’s tests, 1^st^ session; *p* < 0.0001. All other *P’s* > 0.05; **Figure 4A**). This suggests that COC rats attributed more incentive value to cocaine, to the cocaine-paired cues, or both. After extinction sessions, rats received 5 ‘pre-reinstatement’ sessions, where active-lever presses no longer produced cocaine cues. Both groups decreased their active-lever pressing over sessions (Session effect, F_4_, _76_ = 5.77, *p* = 0.0004; **Figure 4B**), and there were no group differences.

**Figure 4.**
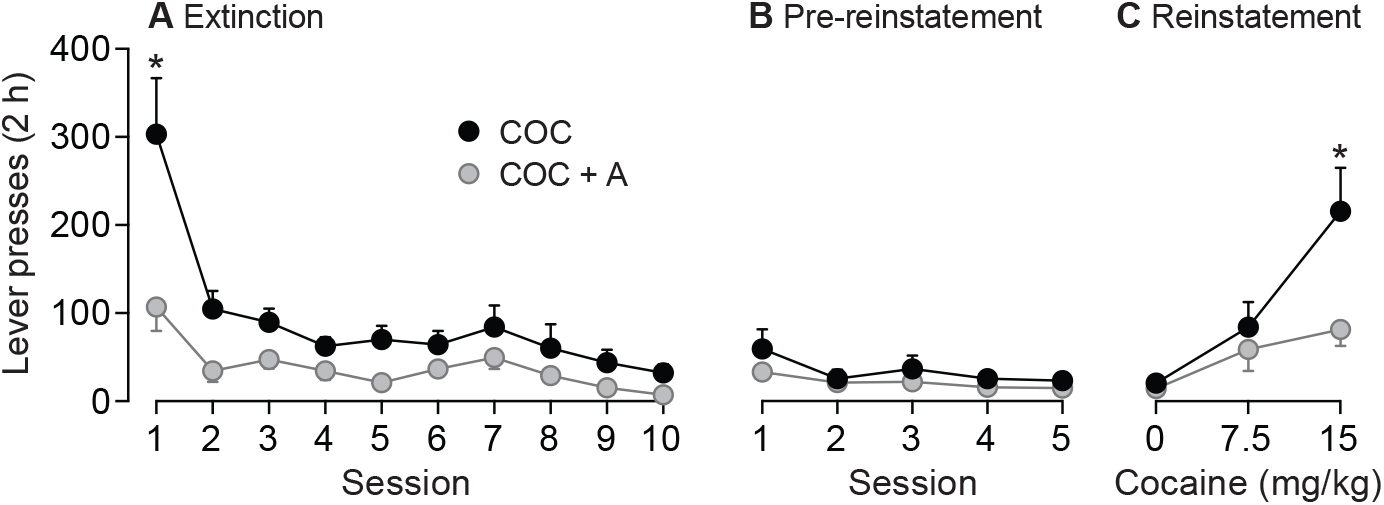
D-amphetamine maintenance during intermittent cocaine self-administration decreased both later responding for cocaine under extinction conditions and cocaine-primed reinstatement of extinguished cocaine-seeking behavior. (**A**) Under extinction conditions, COC + A rats showed less cocaine-seeking behavior than COC rats, but both groups extinguished responding over time. (**B**) During pre-reinstatement sessions (additional extinction sessions, but where cocaine cues were no longer presented) both groups showed very low levels of cocaine-seeking behavior. (**C**) COC + A rats were less vulnerable to cocaine-induced reinstatement of extinguished cocaine-seeking than COC rats. Data are mean ± SEM. n’s = 10-11/group. *P’s < 0.01, COC versus COC + A. COC, Cocaine. A, D-amphetamine.

#### Cocaine-induced reinstatement

Priming injections of cocaine dose-dependently increased active-lever presses in both groups (Dose effect, F_2_, _38_ = 19.26, *p* < 0.0001; **Figure 4C**), but to a greater extent in COC than COC + A rats (Group x Cocaine dose effect, F_2_, _38_ = 5.24, *p* = 0.01; **Figure 4C**), after a 15 mg/kg cocaine prime (Bonferroni’s tests, *p* = 0.003; all other *P’s* > 0.05; **Figure 4C**). Thus, D-amphetamine treatment during intermittent cocaine intake decreased later vulnerability to cocaine-induced relapse, long after the cessation of treatment.

#### Effects of D-amphetamine Maintenance After a History of Intermittent Cocaine Intake

After 14 IntA-sessions (**Figure 2**) and testing under a PR schedule (**Figure 3**), 11 COC rats received 14 additional IntA-sessions (sessions 15-28), now with concomitant D-amphetamine. D-amphetamine was then discontinued and responding for cocaine under a PR schedule was assessed again. Rats escalated their intake over the 28 IntA-sessions (F_27_, _270_ = 3.47, *p* < 0.0001; Bonferroni’s test, IntA-session 1 versus 28, *p* = 0.02; **Figure 5A**). Cocaine-induced locomotion increased over the first 14 IntA-sessions (i.e., without D-amphetamine treatment; Bonferroni’s test; IntA-session 1 versus 14, *p* < 0.0001; **Figure 5B**), and there was no further increment in locomotion over IntA-sessions 15 to 28, when rats were now on D-amphetamine (*p* = 0.66; **Figure 5B**). This is further highlighted in **Figures 5C-D** showing that when locomotion is averaged over the 5-min cocaine periods, there is an increase between IntA-session 1 and 14 (t_10_ = 3.92, *p* = 0.003; **Figure 5C**), but a decrease between IntA-session 15 and 28 (t_10_ = 2.37, *p* = 0.04; **Figure 5D**). Before D-amphetamine treatment, locomotion also followed a spiking pattern during IntA-sessions (**Figures 5E-G**). D-amphetamine attenuated this spiking pattern (**Figures 5H-J**). Thus, D-amphetamine attenuated psychomotor sensitization to cocaine.

**Figure 5.**
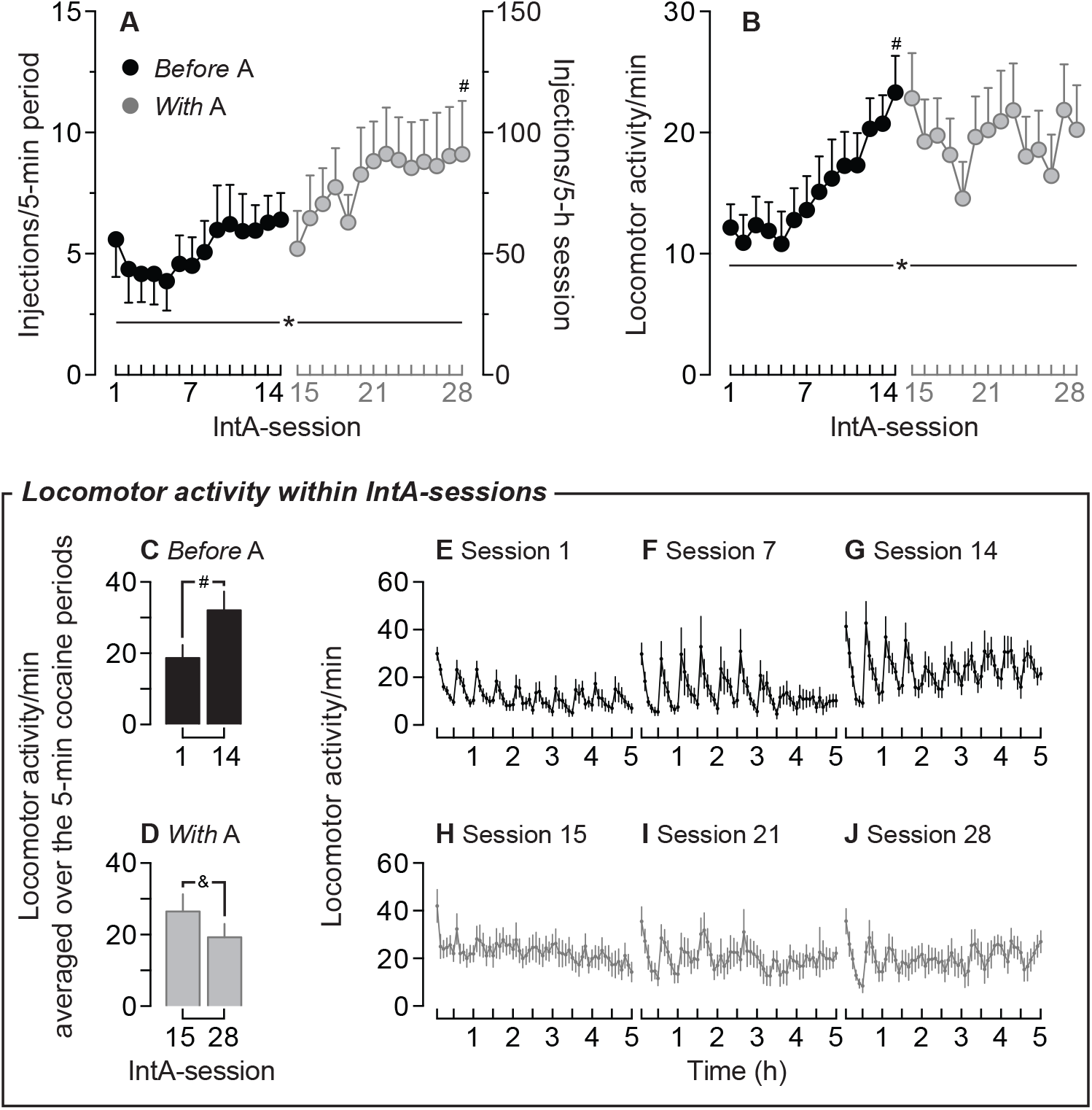
D-amphetamine maintenance does not prevent the escalation of cocaine intake in already cocaine-experienced rats. (**A**) Average number of cocaine injections per 5-min cocaine period (left Y-axis) and total number of cocaine injections per 5-h IntA-session (right Y-axis). Before and with D-amphetamine treatment, rats escalated their cocaine intake over IntA-sessions. (**B**) Locomotor activity increased over intermittent cocaine self-administration sessions, and it stabilized with D-amphetamine. Locomotor activity/min averaged over the 5-minute cocaine periods (**C**) increased from the 1^st^ to the 14^th^ IntA-session (before D-amphetamine) and (**D**) decreased from the 15^th^ to the 28^th^ IntA-session, when rats now received concomitant D-amphetamine treatment. (**E-G**) Locomotor activity during intermittent cocaine self-administration sessions followed a spiking pattern before D-amphetamine, and (**H-J**) D-amphetamine attenuated this spiking effect. Data are mean ± SEM. n = 11. *P’s < 0.0001, Session effect. ^#^P’s < 0.05, versus IntA-session 1. ^&^p < 0.05, versus IntA-session 15. IntA, Intermittent Access. A, D-amphetamine.

#### Responding under a PR schedule

Before and after D-amphetamine treatment, rats responded more for higher cocaine doses under a PR schedule (Dose effect, F_2_, _20_ = 79.03, *p* < 0.0001; **Figure 6A**). Rats earned fewer cocaine injections after D-amphetamine treatment than before (Treatment effect, F_1_, _10_ = 16.51, *p* = 0.002; **Figure 6A**, see also **Figures 6B-D**). After D-amphetamine treatment, cumulative cocaine intake during PR sessions was also decreased (**Figures 6E-G**). Thus, although rats escalated their cocaine intake even further under D-amphetamine (IntA-sessions 15-28, see **Figure 5A**), and they more than doubled their average cumulative cocaine exposure with these additional sessions (average cumulative cocaine intake; IntA-sessions 1-14; 183 mg/kg ± 41 SEM; IntA-sessions 1-28; 463 mg/kg ± 89 SEM), they showed less incentive motivation for cocaine after D-amphetamine treatment. This suggests that while D-amphetamine might not prevent escalation of cocaine intake, it decreases incentive motivation for cocaine. This is unlikely due to repeated testing, because with repeated testing, responding for cocaine under a PR schedule remains stable or even increases [(46), see also (30)].

**Figure 6.**
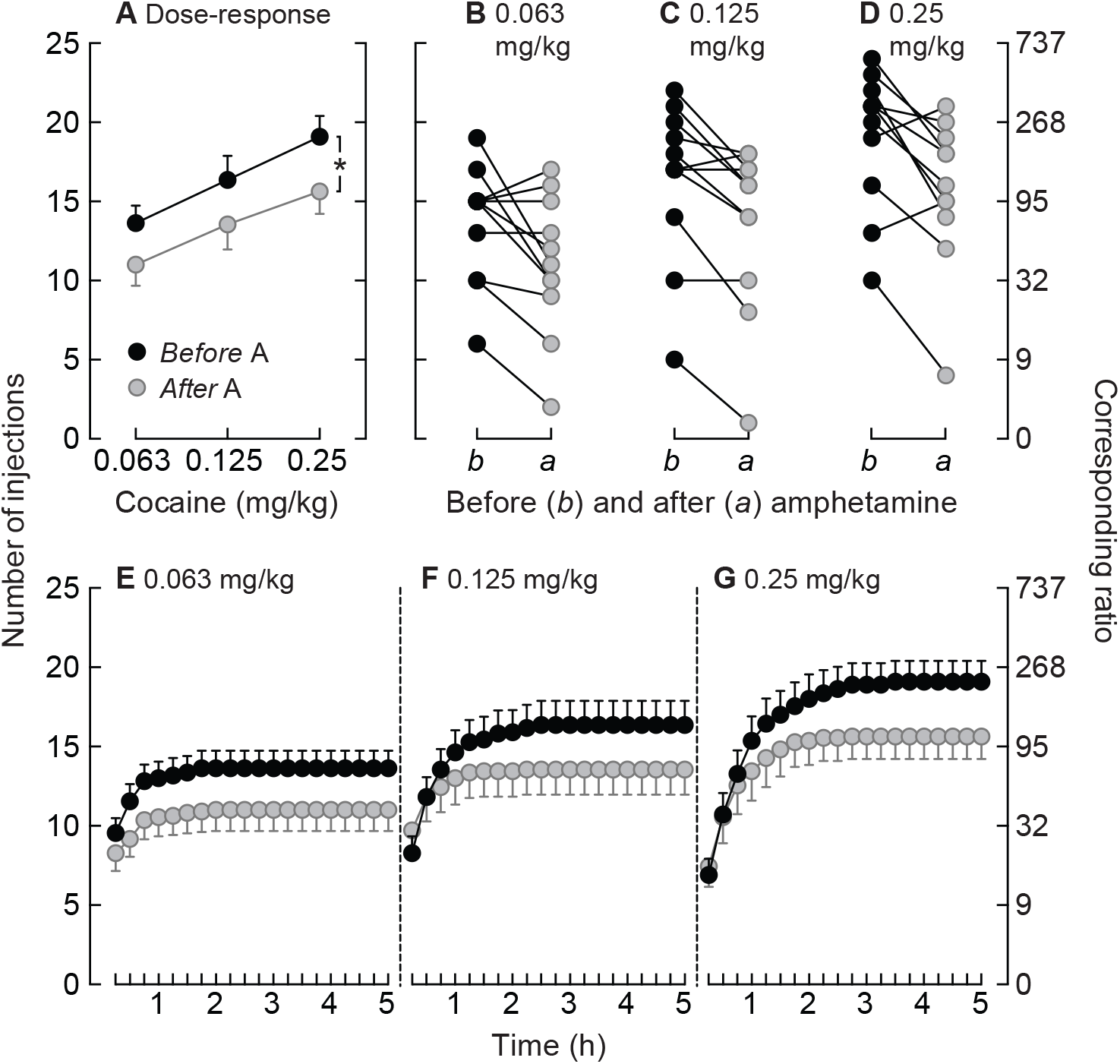
In cocaine-experienced rats, D-amphetamine maintenance during intermittent cocaine self-administration decreases later responding for cocaine under a progressive ratio schedule of reinforcement. (**A**) Number of self-administered cocaine injections (left Y-axis) and corresponding ratio (right Y-axis) before and after D-amphetamine treatment. (**B-D**) Responding for cocaine under progressive ratio in individual rats before (b) and after (a) D-amphetamine treatment. (**E-G**) Cumulative number of self-administered cocaine injections (left Y-axis) and corresponding ratio (right Y-axis) during each 5-h progressive ratio test, as a function of cocaine dose. Data are mean ± SEM. n = 11. *p = 0.002, Treatment effect. A, D-amphetamine.

### Experiment 2: Effects of D-amphetamine Maintenance during Intermittent Cocaine Self-administration on Cocaine’s Potency at the Dopamine Transporter

COC and COC + A rats took similar amounts of cocaine, and D-amphetamine treatment increased saline self-administration (**Figure S2**; unpaired *t*-tests, COC vs. COC + A, t_11_ = 0.55, *p* = 0.59; SAL vs. SAL + A, t_8_ = 2.77, *p* = 0.02). As in experiment 1, locomotion followed a spiking pattern during cocaine self-administration sessions, increased across sessions (**Figure 7A**, black curves), and D-amphetamine blunted both effects. In rats self-administering saline **(Figure 7B**), D-amphetamine initially increased locomotion, but the SAL and SAL + A groups had similar locomotor counts by the last (14^th^) session.

**Figure 7.**
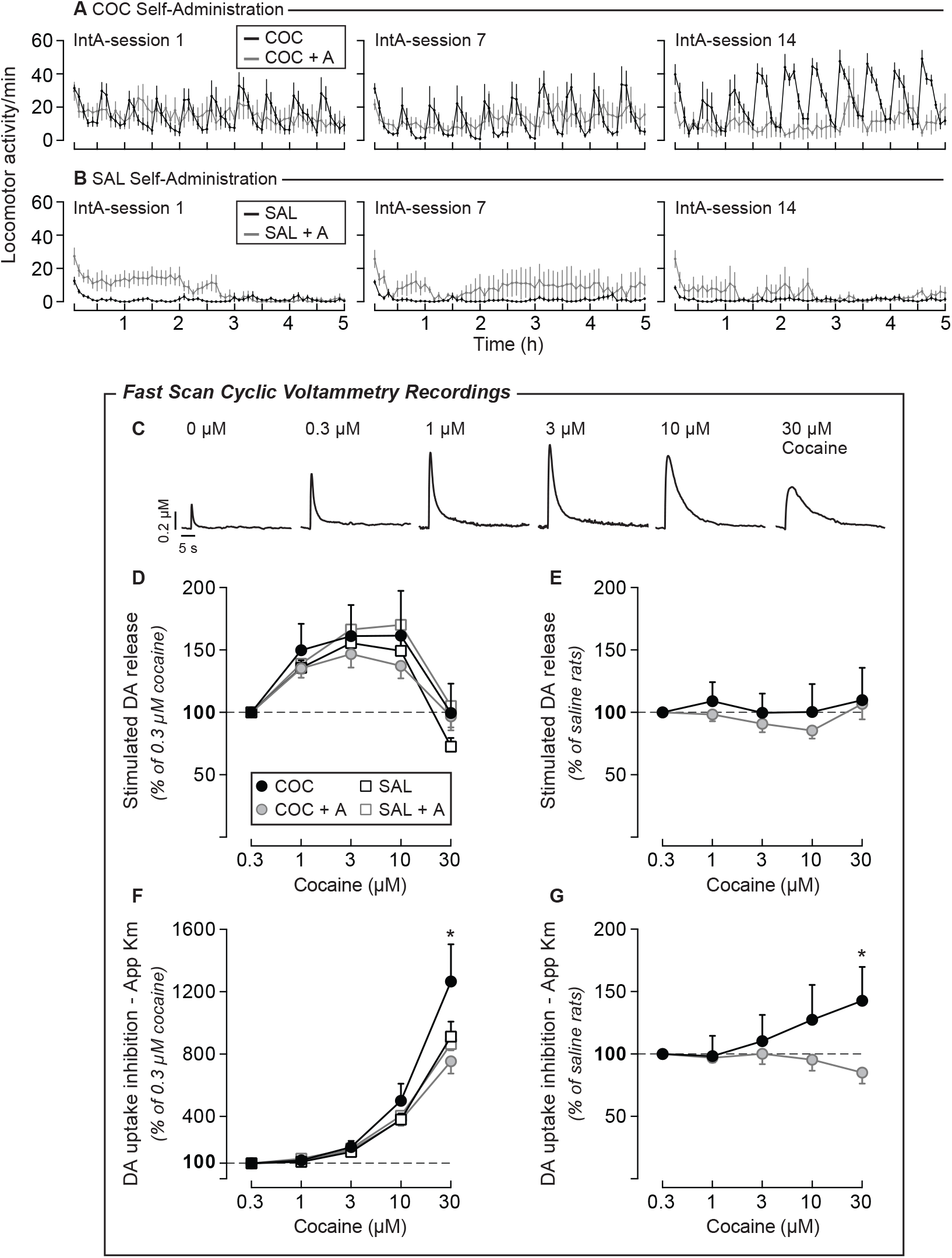
D-amphetamine maintenance during intermittent cocaine intake prevents sensitization to cocaine’s effects at the dopamine transporter. (**A**) As in Experiment 1, intermittent cocaine intake produced spikes in locomotor activity, and spike amplitude increased with more extensive cocaine-taking experience. D-amphetamine attenuated both the locomotion spikes and the increase in spike amplitude over time. (**B**) D-amphetamine increased locomotion in rats self-administering saline. (**C**) Representative traces showing relative extracellular dopamine levels over time evoked by bath-application of different cocaine doses in the nucleus accumbens core. (**D-E**) Electrically-evoked dopamine overflow as a function of cocaine concentration did not change across groups (% Relative to 0.3 μM cocaine in **D** and % relative to SAL rats in **E**). (**F-G**) % of apparent Km as a function of cocaine dose preferentially increased in COC rats and this increase is prevented in COC + A rats (Relative to 0.3 μM cocaine in **F** and relative to SAL rats in **G**). The dose of 30 μM cocaine more effectively inhibited dopamine uptake in COC rats compared to the other groups. Because in **D** and **F**, SAL and SAL + A rats were similar in all measures, they were pooled together in **E** and **G**. Data are mean ± SEM. n = 4-7/group. *p < 0.05, COC versus SAL, COC versus SAL + A and COC versus COC + A in **F-G**. IntA, Intermittent Access. DA, Dopamine. COC, Cocaine. SAL, Saline. A, D-amphetamine.

Five days after cessation of cocaine self-administration/D-amphetamine treatment, we measured cocaine-induced inhibition of DA uptake in the NAcC using *ex vivo* FSCV. **Figure 7C** shows representative FSCV traces following bath-applied cocaine. Increasing cocaine concentrations enhanced stimulated dopamine release, and this did not differ between groups (Percent of 0.3 μM cocaine, Concentration effect, F_4_, _72_ = 36.97, *p* < 0.0001; Group effect, F_3_, _18_ = 0.26, *p* = 0.85; **Figure 7D**; Percent of SAL, Concentration effect, F_4_, _44_ = 1.25, *p* = 0.3; Group effect, F_1_, _11_ = 0.23, *p* = 0.64; **Figure 7E**). As reported previously (33), 30 μM cocaine did not significantly alter stimulated dopamine release. Increasing cocaine concentrations also augmented dopamine uptake inhibition (Concentration effect, F_4, 72_ = 126.7, *p* < 0.0001; **Figure 7F**), and the magnitude of this effect varied as a function of group (Group x Concentration effect, F_12_, _72_ = 2.65, *p* = 0.005; **Figure 7F**; F_4_, _44_ = 3.56, *p* = 0.01; **Figure 7G**). At 30 μM cocaine, dopamine uptake inhibition was greatest in COC rats (Bonferroni’s tests; all *P’s* < 0.05; **Figures 7F-G**). Thus, IntA cocaine experience increased cocaine’s potency at the DAT, as reported previously (29, 33). Importantly, D-amphetamine prevented this neurochemical sensitization. In cocaine-naive, SAL rats, D-amphetamine did not change cocaine-induced dopamine uptake inhibition [D-amphetamine effect, F_1_, _7_ = 0.01, *p* = 0.1; D-amphetamine x Concentration effect, F_4_, _28_ = 0.45, *p* = 0.77, **Figure 7F**; See also (23, 47)]. Thus, D-amphetamine normalizes cocaine potency at the DAT by selectively reversing cocaine-induced neurochemical sensitization, without changing baseline DAT function.

## DISCUSSION

Rats self-administered cocaine on an IntA schedule, with or without D-amphetamine maintenance treatment, and we assessed the development of addiction-like behavior (high motivation for cocaine and drug-induced relapse), psychomotor activity and cocaine-induced inhibition of DA uptake. We report three main findings. First, IntA cocaine self-administration induced psychomotor sensitization, which was attenuated by D-amphetamine treatment, despite no effect on cocaine consumption. Second, in both previously cocaine-naive rats and in cocaine-experienced rats, D-amphetamine treatment decreased incentive motivation for cocaine (measured using PR procedures), responding for cocaine under extinction and cocaine-primed relapse, relative to animals that received cocaine alone. Third, IntA enhanced cocaine’s potency at the DAT, and D-amphetamine prevented this effect, without changing basal DAT function. Thus, D-amphetamine treatment during IntA cocaine self-administration may reduce motivation for cocaine by interacting with the DAT to prevent sensitization-related changes in cocaine potency and dopamine-mediated signalling.

### Long Access versus Intermittent Access

D-amphetamine maintenance therapy can decrease cocaine use in people with addiction (5–8), and so the neurobiological and psychological basis of this effect is of interest. In an earlier study addressing this question Siciliano et al. (2018) trained rats to self-administer cocaine for 6 hrs/day (LgA) (23), and there is now a large literature showing that LgA is especially effective in producing addiction-like behaviors, relative to rats only allowed access for 1-2 hrs/day (Short Access, ShA). These behaviors include escalation of intake, high motivation for drug, a high propensity to relapse and continued drug-seeking in the face of an adverse consequence (24, 48–51). It is also reported that LgA experience produces tolerance in the ability of cocaine to inhibit the DAT (33, 52, 53). Importantly, Siciliano et al. (2018) found that D-amphetamine maintenance treatment not only prevented and reversed the escalation of cocaine intake and the increase in motivation for cocaine otherwise produced by LgA experience (23), but also prevented the tolerance to cocaine’s effect on the DAT produced by LgA (23). They proposed, therefore, that D-amphetamine attenuated addiction-like behavior because it reversed the tolerance produced by extended cocaine use (23). This is consistent with the view that addiction results from tolerance-related adaptations in the DA system, whereby drug-seeking and drug-taking behavior is motivated primarily to overcome this ‘DA deficiency’ and associated anhedonia (54, 55).

However, recent studies using IntA self-administration procedures have begun to paint a very different picture of how cocaine use changes brain and behavior to promote problematic patterns of use [(24–26, 28–33, 56); Reviewed in (19, 20)]. During LgA, brain cocaine concentrations are maintained at a steady high level for the duration of the self-administration session, and the resultant large amount of cocaine consumption was thought to be necessary for the development of addiction-like behavior (48, 57, 58). It turns out this is not the case. Zimmer et al (2011) initially developed the IntA cocaine-self-administration procedure (59), which results in an intermittent, ‘spiking’ pattern in brain cocaine concentrations, because it was thought to better model patterns of cocaine use in humans (27, 60). Importantly, despite much less total cocaine consumption than LgA (comparable to ShA conditions) IntA experience not only also produces escalation of cocaine intake, but is either more effective, or at least as effective, as LgA in producing the addiction-like behaviors described above [(24–26, 28, 30–32, 34); Reviewed in (19, 20)]. Furthermore, rather than producing tolerance, IntA experience enhances (sensitizes) cocaine’s potency in inhibiting DA uptake *ex vivo* (29, 33), and cocaine-induced DA overflow *in vivo* (34). This is consistent with the behavioral and neurobiological effects of experimenter-administered cocaine when given continuously versus intermittently (61, 62).

Given the dramatic differences in the effects of LgA vs. IntA on the DAT, it was important to determine the effects of D-amphetamine treatment on addiction-like behavior and DAT function in rats with IntA experience. D-amphetamine treatment attenuated addiction-like behavior produced by intermittent cocaine intake, as indicated by reductions in motivation for cocaine, responding during extinction and in the magnitude of cocaine-induced reinstatement of cocaine seeking. D-amphetamine was efficacious both in previously cocaine-naive rats and in cocaine-experienced rats, suggesting that D-amphetamine can suppress both the development and the expression of addiction-like behavior. D-amphetamine did not change cocaine’s effects on electrically-evoked dopamine release or cocaine’s potency at the DAT in cocaine-naive rats [See also (23, 47)], but, importantly, it prevented the sensitization of cocaine’s action at the DAT produced by IntA. We hypothesize, therefore, that D-amphetamine reduced the high motivation for cocaine produced by IntA experience by preventing *sensitization* to cocaine’s effects at the DAT, without changing baseline DAT function. This is consistent with an incentive-sensitization view of addiction (63).

### How can both an increase and decrease in DAT function attenuate addiction-like behavior?

It is intriguing that D-amphetamine treatment reduces the development of addiction-like behavior produced by both LgA (23) and IntA cocaine self-administration experience (present findings), but produces apparently opposite effects on DAT function under these two conditions. One possibility is that at least some of the addiction-like behaviors produced by LgA versus IntA experience are due to drug-induced changes in different neuropsychological processes (31). For example, the escalation of intake and the high level of effort expended to obtain cocaine could be because of tolerance to cocaine’s desired effects in the case of LgA, but to sensitization of drug ‘wanting’ in the case of IntA (20, 31, 34). Two lines of evidence support this idea. First, IntA produces sensitization to the psychomotor activating, incentive motivational and dopamine-increasing effects of cocaine (24–26, 28, 29, 31, 33, 34, 42, 43, 56), whereas LgA can decrease the psychomotor activating effects of cocaine and dopamine function, at least soon after the discontinuation of LgA (33, 52, 53, 64). Note that psychomotor sensitization may be expressed after a long period of withdrawal from LgA (50, 56). Second, D-amphetamine treatment reduced the escalation of cocaine intake under LgA conditions (23), but not during IntA (present study), further suggesting that different processes are involved.

We were surprised that D-amphetamine reduced incentive motivation for cocaine, but on average it did not affect the escalation of cocaine intake during IntA-sessions. This was surprising given that D-amphetamine suppressed the development of psychomotor sensitization across IntA-sessions. This could indicate that escalation of cocaine intake during IntA experience does not necessarily reflect sensitization of an appetitive process, and that cocaine consumption and appetitive responding for the drug are dissociable (24, 26, 43–45). Indeed, we found that cocaine intake during IntA-sessions did not predict responding for the drug under a PR schedule [see also (26, 28, 42)]. Alternatively, perhaps D-amphetamine’s therapeutic effects are most robust after the cessation of treatment, because rats can take enough cocaine during IntA-sessions to overwhelm any effect of ongoing D-amphetamine treatment on the DAT. The implication is that D-amphetamine maintenance reduces incentive motivation for cocaine by inoculating against cocaine-induced, sensitization-related plasticity at the DAT. Future work should also examine behavioral and neurochemical outcome at different timepoints after the discontinuation of continuous D-amphetamine, as the effects of continuous psychostimulant drug treatment can change following longer withdrawal periods (50, 56, 65).

IntA not only produces successive ‘spikes’ in brain cocaine levels (24, 59) but in psychomotor activity, as seen here. Typically, there is a strong correlation between brain concentrations of cocaine, extracellular concentrations of DA in the striatum, and the time course of the psychomotor activating effects of cocaine (66–68). It is interesting, therefore, that D-amphetamine ‘flattened’ the spikes in psychomotor activity otherwise produced by intermittent cocaine intake [see also (28)]. We speculate that D-amphetamine may have also attenuated the intermittent ‘spikes’ in DA that would otherwise occur with IntA self-administration. The importance of intermittency in producing sensitization-related changes in brain and behavior has been long-recognised (69–73), and therefore, D-amphetamine may have therapeutic effects because it blunts the intermittent spikes in striatal dopamine activity that promote the induction of sensitization. This hypothesis remains to be tested.

## CONCLUSIONS

Using a cocaine self-administration procedure that models intermittent cocaine taking in humans (24, 27, 59), we report that intermittent cocaine use produces robust sensitization to both the psychomotor-activating and DAT-inhibiting effects of cocaine. This extends reports that intermittent-access cocaine promotes sensitization to cocaine’s dopamine-elevating effects [(29, 33, 34); See (20) for review]. The original finding here is that D-amphetamine attenuates the cocaine-taking and -seeking behaviors otherwise promoted by intermittent cocaine intake, and that this is associated with blunting of sensitization-related changes in cocaine potency at the DAT. This is consistent with the view that the transition to cocaine addiction involves sensitization-related neuroplasticity (63), and therefore, treatments that reverse this may be especially efficacious.

## Acknowledgments and author contributions

This research was supported by the Canada Foundation for Innovation and the Canadian Institutes of Health Research to ANS (grant numbers 24326 and 157572, respectively). ANS is supported by a salary grant from the Fonds de Recherche du Québec – Santé (grant number 28988). FA is supported by a PhD fellowship from the Groupe de Recherche sur le Système Nerveux Central; Research in the LET lab was supported by a grant from the National Science and Engineering Research Council of Canada (grant 05230). We are thankful to Dr. David CS Roberts for generous advice on administering D-amphetamine treatment during cocaine self-administration. We thank Dr. Erin S Calipari for advice on FSCV and Marie-Josée Bourque for her help in optimizing FSCV electrode construction. We thank Dr Kent C Berridge whose work has inspired interpretation of our findings.

FA designed the research, performed all behavioral experiments and analyzed all data with guidance from ANS. MPB contributed to rat behavioral testing in Experiment 1. BDL performed FSCV recordings with guidance from LET. VJ extracted apparent Km values from FSCV data. FA, TER and ANS wrote the article with revisions from BDL, MPB, VJ and LET.

## SUPPLEMENTAL INFORMATION

### SUPPLEMENTAL METHODS

#### Subjects

Male Wistar rats (200-250 g; Charles River Laboratories, St Constant, Qc) were housed 1/cage under a 12-h reverse light-dark cycle (lights off at 08:30 am). Upon arrival to the animal colony, rats were left undisturbed for 3 days with ad libitum access to food and water. After this, rats were handled daily during the dark phase of the circadian cycle and food was restricted (74) to 25 g/day with the following exceptions: 15 g/day during food self-administration training and 35 g on the days surrounding catheter implantation. Water was available *ad libitum*. The animal care committee of the Université de Montréal approved all procedures involving animals.

#### Food self-administration training

All self-administration training and testing took place in standard operant conditioning cages that also contained infrared photocells to measure horizontal locomotor activity. At the beginning of each session, the house-light was illuminated, and the levers were inserted. Pressing the active lever was reinforced with one banana-flavored food pellet (45 mg; grain-based; VWR, Montreal, QC) under a fixed ratio 1 schedule (FR1). Upon each reward delivery, the light above the active lever was turned on and both levers were retracted during a 20-s timeout period. After this timeout period, the light above the active lever was turned off and both levers were again inserted into the cage. Pressing the inactive lever had no programmed consequences. Each food training session lasted 1 h or until 100 food pellets were delivered. At the end of each session, the house-light was turned off and the levers were retracted. All rats met the acquisition criterion (taking ~100 pellets/session on two consecutive days) in 2-3 days. Next, for two sessions, food pellets were available under FR3. The i.v. catheters were then implanted into the jugular vein of the rats.

#### Catheter implantation

On the day following the end of food training, catheters were implanted into the jugular vein of the rats (75–77), under isofluorane anesthesia (5% for induction, 2-3% for maintenance). Homemade catheters consisted of a 12.5-cm length of silastic tubing linked to a stainless-steel cannula (C313G-5UP). Dental cement was then used to affix the cannula to a circular piece of nylon mesh. Two silicone bubbles were placed on the silastic tubing. These bubbles served to fasten the catheter to the jugular vein and to the chest muscle with suture thread. The cannula with the nylon mesh was set to lie subcutaneously in between the rats’ scapulae. On the day of surgery, rats received an intramuscular injection of a penicillin antibiotic (Procillin; CDMV, St Hyacinthe, Qc) and a subcutaneous injection of the anti-inflammatory agent Carprofen (Rimadyl; CDMV, St Hyacinthe, Qc). To avoid coagulation, catheters were flushed each day with a sterile saline solution and every other day with a heparinized saline solution (0.2 mg/ml; Sigma-Aldrich, Oakville, ON).

#### Drugs

Cocaine hydrochloride (Medisca Pharmaceutique, St Laurent, Qc; 0.25 mg/kg/injection, delivered over 5 s) and D-amphetamine (A; Sigma-Aldrich, Dorset, UK) were dissolved in 0.9% saline. An osmotic minipump (Alzet model 2ML2, Durect, Cupertino, CA, USA) set to deliver 5 mg/kg/day D-amphetamine for 14 days was implanted in each COC + A or SAL + A rat. This dose and regimen of D-amphetamine administration was selected based on prior work in rats (14, 15) showing that it decreases responding for cocaine under a progressive ratio schedule of reinforcement, but it increases responding for sucrose pellets and does not reduce body weight, suggesting no debilitating or overtly toxic effects. Of note, D-amphetamine administration via minipump can induce neurotoxicity in rats, as indicated by degenerating axon terminals or tyrosine hydroxylase immunoreactive patches, but at doses above 16-20 mg/kg/day (78–80). We weighed the rats before minipump implantation and we then estimated body weight mid-way through the projected 14-day D-amphetamine treatment. Actual body weights on day 7 of D-amphetamine treatment showed that rats received ~5.1 mg/kg/day D-amphetamine.

#### Cocaine self-administration training

The rats were tested in operant conditioning cages (Med Associates, St Albans, VT) containing two retractable levers, a cue light above each lever and infrared photocells to measure horizontal locomotor activity. After acquisition of operant responding for food pellets, rats learned to self-administer cocaine (0.25 mg/kg/injection, delivered over 5 s) during 1-h sessions (1 session/day) under a fixed ratio 3 schedule of reinforcement (FR3). Each session started with illumination of the house light and insertion of two levers. Pressing the active lever produced a cocaine injection, illumination of a light cue above the active lever and retraction of both levers for a 20-s timeout period during which lever presses were not reinforced. The light cue was on for the 5-s injection and the ensuing 20-s time out period. Pressing the inactive lever had no consequences. We considered that rats acquired reliable cocaine self-administration behavior when on two consecutive days, they took ≥ 6 injections/session, at regular intervals throughout the session, and pressed ≥ 2 times more on the active versus inactive lever (26, 28, 42).

#### Osmotic minipump implantation

After acquiring cocaine self-administration, rats were anaesthetized with isofluorane and a minipump was inserted subcutaneously on the animals’ backs, caudal to the catheter. Rats allocated to the D-amphetamine condition were implanted with a minipump filled with D-amphetamine. These are the COC + A rats in Experiment 1, and COC + A as well as SAL + A rats in Experiment 2. In Experiment 1, rats allocated to the COC group were implanted with a minipump filled with saline or received a sham surgery consisting of an incision and sutures. In Experiment 2, COC and SAL rats received a sham surgery.

#### Responding under a PR schedule

Obtaining each cocaine injection (0.063-0.25 mg/kg/injection, one session/dose, counterbalanced) required an exponentially increasing number of lever presses, according to the following formula: [5 × *e*^(*number of injection* × 0.2)^ − 5] (35). Sessions ended after 5 hours or when 1 h elapsed without a self-administered injection.

#### Extinction

For 10, 2-h sessions, rats were tethered to the infusion apparatus and had continuous lever access. Each three active-lever presses produced exteroceptive cocaine-associated cues (i.e., syringe pump activation, lever retraction and illumination of the cue light for 5 seconds), but no cocaine. The day following the last extinction session, rats received five, 2-h pre-reinstatement sessions to further decrease the influence of cocaine cues on subsequent reinstatement testing (36–38). During these sessions, the rats were placed in the operant cages but were not connected to the infusion apparatus. Cages were also unlit and lever presses had no consequences.

#### Effects of cocaine on dopamine uptake inhibition in the nucleus accumbens core (NAcC) ex vivo

We used a fast scan cyclic voltammetry (FSCV) protocol similar to that described in our previous work (81–83). Rats were injected i.p. with sodium pentobarbital (90 mg/kg) and perfused with a NMDG-based solution (in mM; 92 NaCl, 2.5 KCl, 1.25 NaH_2_PO_4_, 30 NaHCO_3_, 20 HEPES, 25 glucose, 2 thiourea, 5 ascorbic acid, 3 sodium pyruvate, 2 CaCl_2_, 2 MgSO_4_) for at least 1 hour at room temperature. The brain was extracted, and 300-μm-thick coronal slices were prepared in ice-cold (0 to 4°C) NMDG solution using a vibrating blade microtome (Leica VT1000S). Slices were first placed in a 32°C NMDG solution for 12 min and then in an oxygenated HEPES-buffered resting solution (in mM; 92 NaCl, 2.5 KCl, 1.25 NaH_2_PO_4_, 30 NaHCO_3_, 20 HEPES, 25 glucose, 2 thiourea, 5 ascorbic acid, 3 sodium pyruvate, 2 CaCl_2_, 2 MgSO_4_) for at least 1 h at room temperature. A slice containing the NAcC [approximately +1.7 mm from Bregma (84)] was placed in the recording chamber and continuously perfused (1 ml/min) with oxygenated aCSF (in mM; 119 NaCl, 2.5 KCl, 1.25 NaH_2_PO_4_, 24 NaHCO_3_, 12.5 glucose, 2.4 CaCl_2_, 1.3 MgSO_4_) at 32°C. A carbon-fiber electrode (CFE, ~7 μm in diameter) was placed ~100 μm below the surface of the slice, into the NAcC and centered between the two poles of the bipolar stimulating electrode (Plastics One, Roanoke, VA, US) placed on the slice surface. Single-pulse electrical stimulations (400 μA; 1 ms) were generated every 5 minutes to evoke dopamine release and the potential at the CFE was scanned according to a 10-ms triangular voltage wave (−0.4 to 1 to −0.4 V vs. Ag/AgCl, at the rate of 300 V/s). Dopamine release was analyzed as the peak height of electrically-evoked extracellular dopamine overflow. Once three stable responses were recorded, increasing concentrations of cocaine (0.3, 1, 3, 10, 30 μM) were cumulatively applied to the bath. Once 3 stable responses were recorded at a given cocaine concentration, the next concentration was applied.

#### Analysis of the kinetics of fast-scan cyclic voltammetry data

Dopamine reuptake was modelled from the rate of recovery of the dopamine signal using Michaelis-Menten kinetics (85). First, nonlinear least-square optimization was applied to fit a three-parameter exponential function with baseline shift to the reuptake phase of the dopamine response. The time constant τ (tau) of the exponential corresponds to the half-life divided by log(2) and is related to the Michaelis-Menten parameters Vmax and Km. It can be shown by integration that the area under the exponential function and the area under an equivalent Michaelis-Menten curve are equal if and only if τ. Vmax = Km + [DA]/2, where [DA] is dopamine peak height. Under the assumptions that Km = 0.18 μM in cocaine-free conditions and Vmax remains constant during each experiment (33), this formula provides an estimate of Km (apparent Km, or app. Km) based on τ and [DA], while avoiding overfitting. The parameters [DA] and app. Km were extracted from each recording. In cocaine-free conditions, signals were of relatively low amplitude, such that signal-to-noise ratio was often suboptimal for accurate parameter identification. As such, we computed kinetic parameters as percentages relative to 0.3 μM cocaine. Note that in Calipari et al (2013), app. Km at 0.3 μM cocaine was only < 10-20% greater than control levels (see their Figure 3C), which remains small compared to the increases evoked by higher cocaine concentrations (> 1000%).

#### Statistical analyses

Two-way repeated measures (RM) ANOVAs (Group x Session/Time in min/Dose, Session and Dose as within-subjects variables) were used to analyze cocaine intake and locomotor activity over IntA-sessions, and cocaine intake over PR sessions. Three-way RM ANOVAs (Group x Session x Time in min; Session and Time as within-subjects variables) were used to analyse locomotor activity within IntA-sessions. When comparing COC to COC + A rats, ‘Session’, ‘Time in min’ and ‘Dose’ were analyzed as within-subjects variables and Group as a between-subjects variable. Two-way RM ANOVA was used to analyze lever-presses during extinction, pre-reinstatement (Group x Session; the latter as a within-subjects variable) and reinstatement sessions (Group x Cocaine dose, the latter as a within-subjects variable). When comparing responding before and with D-amphetamine treatment in the same rats, one-way RM ANOVA was used to analyze cocaine intake and locomotor activity over the 28 IntA-sessions. Locomotor activity averaged over the 5-min cocaine periods was analyzed before and with D-amphetamine treatment using paired *t*-tests (IntA-session 1 versus 14 and IntA-session 15 versus 28). Two-way RM ANOVA (Group x Dose, both as within-subjects variables) was used to compare responding under PR before and after D-amphetamine. Finally, two-way RM ANOVA was used to analyze group differences in dopamine overflow and apparent Km as a function of cocaine concentration (0.3 to 30 μM; Group x Cocaine concentration; the latter as a within-subjects variable).

### SUPPLEMENTAL FIGURES

**Figure S1.**
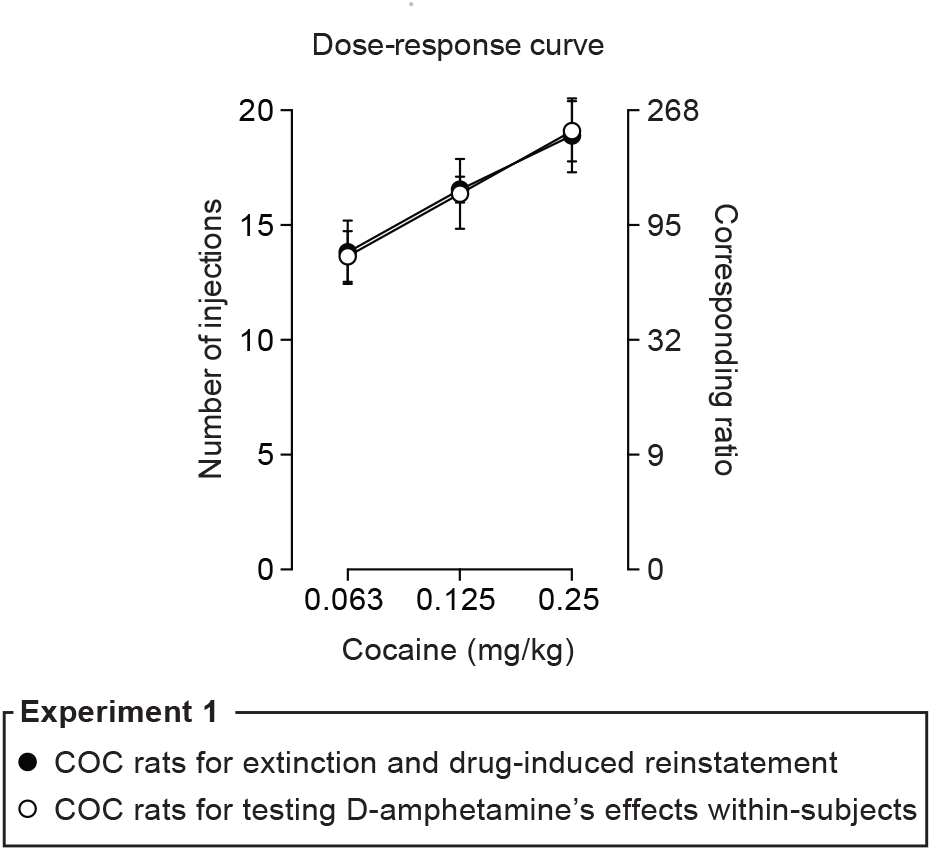
Responding for cocaine under a progressive ratio schedule of reinforcement in COC rats used in Experiment 1. In Experiment 1, after being tested for responding for cocaine (0.063-0.25 mg/kg/infusion) under a progressive ratio schedule of reinforcement, the 22 COC rats were divided into two groups. One group was then tested for extinction and cocaine-induced reinstatement of cocaine-seeking behavior. The other group was given additional IntA-sessions to self-administer cocaine, but now with D-amphetamine treatment. The figure shows that motivation to take cocaine was comparable in the two groups before this division (Dose effect, F_2, 40_ = 36.23, p < 0.0001; Group effect, F_1_, _20_ = 0.001, p = 0.97; Dose x Group effect, F_2_, _40_ = 0.06, p = 0.94). Data are mean ± SEM. n = 11/group. IntA, Intermittent Access. COC, Cocaine. A, D-amphetamine

**Figure S2.**
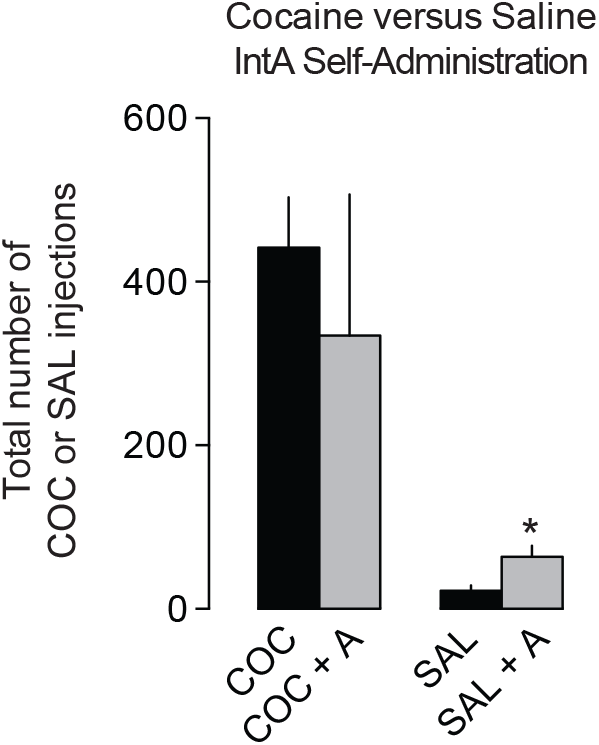
D-amphetamine maintenance did not significantly change total cocaine over the 14 intermittent-access self-administration sessions. D-amphetamine increased inter-individual variability in cocaine intake, but average COC intake was equivalent with or without D-amphetamine maintenance (t_11_ = 0.55, p = 0.59). D-amphetamine maintenance increased the number of self-administered saline injections (t_8_ = 2.77, *p = 0.02, SAL versus SAL + A). IntA, Intermittent Access. COC, Cocaine. SAL, Saline. A, D-amphetamine.

## REFERENCES

1. Vocci F, Ling W (2005): Medications development: successes and challenges. Pharmacol Ther. 108:94–108.

2. Fischer B, Blanken P, Da Silveira D, Gallassi A, Goldner EM, Rehm J, et al. (2015): Effectiveness of secondary prevention and treatment interventions for crack-cocaine abuse: a comprehensive narrative overview of English-language studies. Int J Drug Policy. 26:352–363.

3. Sanchez-Hervas E (2016): Cocaine addiction: treatments and future perspectives. Trends Psychiatry Psychother. 38:242–243.

4. Negus SS, Henningfield J (2015): Agonist Medications for the Treatment of Cocaine Use Disorder. Neuropsychopharmacology. 40:1815–1825.

5. Grabowski J, Rhoades H, Schmitz J, Stotts A, Daruzska LA, Creson D, et al. (2001): Dextroamphetamine for cocaine-dependence treatment: a double-blind randomized clinical trial. J Clin Psychopharmacol. 21:522–526.

6. Greenwald MK, Lundahl LH, Steinmiller CL (2010): Sustained release d-amphetamine reduces cocaine but not ‘speedball’-seeking in buprenorphine-maintained volunteers: a test of dual-agonist pharmacotherapy for cocaine/heroin polydrug abusers. Neuropsychopharmacology. 35:2624–2637.

7. Rush CR, Stoops WW, Sevak RJ, Hays LR (2010): Cocaine choice in humans during D-amphetamine maintenance. J Clin Psychopharmacol. 30:152–159.

8. Shearer J, Wodak A, van Beek I, Mattick RP, Lewis J (2003): Pilot randomized double blind placebo-controlled study of dexamphetamine for cocaine dependence. Addiction. 98:1137–1141.

9. Czoty PW, Gould RW, Martelle JL, Nader MA (2011): Prolonged attenuation of the reinforcing strength of cocaine by chronic d-amphetamine in rhesus monkeys. Neuropsychopharmacology. 36:539–547.

10. Czoty PW, Martelle JL, Nader MA (2010): Effects of chronic d-amphetamine administration on the reinforcing strength of cocaine in rhesus monkeys. Psychopharmacology (Berl). 209:375–382.

11. Negus SS (2003): Rapid assessment of choice between cocaine and food in rhesus monkeys: effects of environmental manipulations and treatment with d-amphetamine and flupenthixol. Neuropsychopharmacology. 28:919–931.

12. Negus SS, Mello NK (2003): Effects of chronic d-amphetamine treatment on cocaine- and food-maintained responding under a second-order schedule in rhesus monkeys. Drug Alcohol Depend. 70:39–52.

13. Negus SS, Mello NK (2003): Effects of chronic d-amphetamine treatment on cocaine- and food-maintained responding under a progressive-ratio schedule in rhesus monkeys. Psychopharmacology (Berl). 167:324–332.

14. Chiodo KA, Lack CM, Roberts DC (2008): Cocaine self-administration reinforced on a progressive ratio schedule decreases with continuous D-amphetamine treatment in rats. Psychopharmacology (Berl). 200:465–473.

15. Chiodo KA, Roberts DC (2009): Decreased reinforcing effects of cocaine following 2 weeks of continuous D-amphetamine treatment in rats. Psychopharmacology (Berl). 206:447–456.

16. Thomsen M, Barrett AC, Negus SS, Caine SB (2013): Cocaine versus food choice procedure in rats: environmental manipulations and effects of amphetamine. J Exp Anal Behav. 99:211–233.

17. Zimmer BA, Chiodo KA, Roberts DC (2014): Reduction of the reinforcing effectiveness of cocaine by continuous D-amphetamine treatment in rats: importance of active self-administration during treatment period. Psychopharmacology (Berl). 231:949–954.

18. Ritz MC, Lamb RJ, Goldberg SR, Kuhar MJ (1987): Cocaine receptors on dopamine transporters are related to self-administration of cocaine. Science. 237:1219–1223.

19. Allain F, Minogianis EA, Roberts DC, Samaha AN (2015): How fast and how often: The pharmacokinetics of drug use are decisive in addiction. Neurosci Biobehav Rev. 56:166–179.

20. Kawa AB, Allain F, Robinson TE, Samaha AN (2019): The transition to cocaine addiction: the importance of pharmacokinetics for preclinical models. Psychopharmacology (Berl). 236:1145–1157.

21. Lile JA (2006): Pharmacological determinants of the reinforcing effects of psychostimulants: Relation to agonist substitution treatment. Exp Clin Psychopharmacol. 14:20–33.

22. Samaha AN, Robinson TE (2005): Why does the rapid delivery of drugs to the brain promote addiction? Trends Pharmacol Sci. 26:82–87.

23. Siciliano CA, Saha K, Calipari ES, Fordahl SC, Chen R, Khoshbouei H, et al. (2018): Amphetamine reverses escalated cocaine intake via restoration of dopamine transporter conformation. J Neurosci. 38:484–497.

24. Zimmer BA, Oleson EB, Roberts DC (2012): The motivation to self-administer is increased after a history of spiking brain levels of cocaine. Neuropsychopharmacology. 37:1901–1910.

25. Algallal H, Allain F, Ndiaye NA, Samaha AN (2019): Sex differences in cocaine self-administration behaviour under long access versus intermittent access conditions. Addict Biol.e12809.

26. Allain F, Bouayad-Gervais K, Samaha AN (2018): High and escalating levels of cocaine intake are dissociable from subsequent incentive motivation for the drug in rats. Psychopharmacology (Berl). 235:317–328.

27. Beveridge TJR, Wray P, Brewer A, Shapiro B, Mahoney JJ, Newton TF (2012): Analyzing human cocaine use patterns to inform animal addiction model development. Published abstract for the College on Problems of Drug Dependence Annual Meeting, Palm Springs, CA.

28. Allain F, Samaha AN (2018): Revisiting long-access versus short-access cocaine self-administration in rats: intermittent intake promotes addiction symptoms independent of session length. Addict Biol. 24:641–651.

29. Calipari ES, Siciliano CA, Zimmer BA, Jones SR (2015): Brief intermittent cocaine self-administration and abstinence sensitizes cocaine effects on the dopamine transporter and increases drug seeking. Neuropsychopharmacology. 40:728–735.

30. James MH, Stopper CM, Zimmer BA, Koll NE, Bowrey HE, Aston-Jones G (2019): Increased Number and Activity of a Lateral Subpopulation of Hypothalamic Orexin/Hypocretin Neurons Underlies the Expression of an Addicted State in Rats. Biol Psychiatry. 85:925–935.

31. Kawa AB, Bentzley BS, Robinson TE (2016): Less is more: prolonged intermittent access cocaine self-administration produces incentive-sensitization and addiction-like behavior. Psychopharmacology (Berl). 233:3587–3602.

32. Nicolas C, Russell TI, Pierce AF, Maldera S, Holley A, You ZB, et al. (2019): Incubation of Cocaine Craving After Intermittent-Access Self-administration: Sex Differences and Estrous Cycle. Biol Psychiatry. 85:915–924.

33. Calipari ES, Ferris MJ, Zimmer BA, Roberts DC, Jones SR (2013): Temporal pattern of cocaine intake determines tolerance vs sensitization of cocaine effects at the dopamine transporter. Neuropsychopharmacology. 38:2385–2392.

34. Kawa AB, Valenta AC, Kennedy RT, Robinson TE (2019): Incentive and dopamine sensitization produced by intermittent but not long access cocaine self-administration. Eur J Neurosci. 50:2663–2682.

35. Richardson NR, Roberts DC (1996): Progressive ratio schedules in drug self-administration studies in rats: a method to evaluate reinforcing efficacy. J Neurosci Methods. 66:1–11.

36. De Vries TJ, Schoffelmeer AN, Binnekade R, Mulder AH, Vanderschuren LJ (1998): Drug-induced reinstatement of heroin- and cocaine-seeking behaviour following long-term extinction is associated with expression of behavioural sensitization. Eur J Neurosci. 10:3565–3571.

37. Wakabayashi KT, Weiss MJ, Pickup KN, Robinson TE (2010): Rats markedly escalate their intake and show a persistent susceptibility to reinstatement only when cocaine is injected rapidly. J Neurosci. 30:11346–11355.

38. Deroche-Gamonet V, Piat F, Le Moal M, Piazza PV (2002): Influence of cue-conditioning on acquisition, maintenance and relapse of cocaine intravenous self-administration. Eur J Neurosci. 15:1363–1370.

39. Dong Y, Taylor JR, Wolf ME, Shaham Y (2017): Circuit and Synaptic Plasticity Mechanisms of Drug Relapse. The Journal of Neuroscience. 37:10867.

40. Stewart J, de Wit H (1987): Reinstatement of Drug-Taking Behavior as a Method of Assessing Incentive Motivational Properties of Drugs. In: Bozarth MA, editor. Methods of Assessing the Reinforcing Properties of Abused Drugs. New York, NY: Springer New York, pp 211–227.

41. Pitchers KK, Wood TR, Skrzynski CJ, Robinson TE, Sarter M (2017): The ability for cocaine and cocaine-associated cues to compete for attention. Behav Brain Res. 320:302–315.

42. Allain F, Roberts DC, Levesque D, Samaha AN (2017): Intermittent intake of rapid cocaine injections promotes robust psychomotor sensitization, increased incentive motivation for the drug and mGlu2/3 receptor dysregulation. Neuropharmacology. 117:227–237.

43. Kawa AB, Robinson TE (2019): Sex differences in incentive-sensitization produced by intermittent access cocaine self-administration. Psychopharmacology (Berl). 236:625–639.

44. Nicola SM, Deadwyler SA (2000): Firing rate of nucleus accumbens neurons is dopamine-dependent and reflects the timing of cocaine-seeking behavior in rats on a progressive ratio schedule of reinforcement. J Neurosci. 20:5526–5537.

45. Oleson EB, Roberts DC (2009): Behavioral economic assessment of price and cocaine consumption following self-administration histories that produce escalation of either final ratios or intake. Neuropsychopharmacology. 34:796–804.

46. Liu Y, Roberts DC, Morgan D (2005): Sensitization of the reinforcing effects of self-administered cocaine in rats: effects of dose and intravenous injection speed. Eur J Neurosci. 22:195–200.

47. Davidson C, Lee TH, Ellinwood EH (2005): Acute and chronic continuous methamphetamine have different long-term behavioral and neurochemical consequences. Neurochem Int. 46:189–203.

48. Ahmed SH, Koob GF (1998): Transition from moderate to excessive drug intake: change in hedonic set point. Science. 282:298–300.

49. Paterson NE, Markou A (2003): Increased motivation for self-administered cocaine after escalated cocaine intake. Neuroreport. 14:2229–2232.

50. Ferrario CR, Gorny G, Crombag HS, Li Y, Kolb B, Robinson TE (2005): Neural and behavioral plasticity associated with the transition from controlled to escalated cocaine use. Biol Psychiatry. 58:751–759.

51. Xue Y, Steketee JD, Sun W (2012): Inactivation of the central nucleus of the amygdala reduces the effect of punishment on cocaine self-administration in rats. Eur J Neurosci. 35:775–783.

52. Ferris MJ, Mateo Y, Roberts DC, Jones SR (2011): Cocaine-insensitive dopamine transporters with intact substrate transport produced by self-administration. Biol Psychiatry. 69:201–207.

53. Calipari ES, Ferris MJ, Jones SR (2014): Extended access of cocaine self-administration results in tolerance to the dopamine-elevating and locomotor-stimulating effects of cocaine. J Neurochem. 128:224–232.

54. Koob GF, Volkow ND (2016): Neurobiology of addiction: a neurocircuitry analysis. Lancet Psychiatry. 3:760–773.

55. Volkow ND, Koob GF, McLellan AT (2016): Neurobiologic Advances from the Brain Disease Model of Addiction. N Engl J Med. 374:363–371.

56. Carr CC, Ferrario CR, Robinson TE (2019): Intermittent access cocaine self-administration produces psychomotor sensitization: effects of withdrawal, sex and cross-sensitization. bioRxiv.859520.

57. Ahmed SH (2012): The science of making drug-addicted animals. Neuroscience. 211:107–125.

58. Edwards S, Koob GF (2013): Escalation of drug self-administration as a hallmark of persistent addiction liability. Behav Pharmacol. 24:356–362.

59. Zimmer BA, Dobrin CV, Roberts DC (2011): Brain-cocaine concentrations determine the dose self-administered by rats on a novel behaviorally dependent dosing schedule. Neuropsychopharmacology. 36:2741–2749.

60. Ward AS, Haney M, Fischman MW, Foltin RW (1997): Binge cocaine self-administration in humans: intravenous cocaine. Psychopharmacology (Berl). 132:375–381.

61. Izenwasser S, Cox BM (1990): Daily cocaine treatment produces a persistent reduction of [3H]dopamine uptake in vitro in rat nucleus accumbens but not in striatum. Brain Res. 531:338–341.

62. Izenwasser S, Cox BM (1992): Inhibition of dopamine uptake by cocaine and nicotine: tolerance to chronic treatments. Brain Res. 573:119–125.

63. Robinson TE, Berridge KC (1993): The neural basis of drug craving: an incentive-sensitization theory of addiction. Brain Res Brain Res Rev. 18:247–291.

64. Willuhn I, Burgeno LM, Groblewski PA, Phillips PE (2014): Excessive cocaine use results from decreased phasic dopamine signaling in the striatum. Nat Neurosci. 17:704–709.

65. Dalia AD, Norman MK, Tabet MR, Schlueter KT, Tsibulsky VL, Norman AB (1998): Transient amelioration of the sensitization of cocaine-induced behaviors in rats by the induction of tolerance. Brain Res. 797:29–34.

66. Minogianis EA, Shams WM, Mabrouk OS, Wong JMT, Brake WG, Kennedy RT, et al. (2019): Varying the rate of intravenous cocaine infusion influences the temporal dynamics of both drug and dopamine concentrations in the striatum. Eur J Neurosci. 50:2054–2064.

67. Shou M, Ferrario CR, Schultz KN, Robinson TE, Kennedy RT (2006): Monitoring dopamine in vivo by microdialysis sampling and on-line CE-laser-induced fluorescence. Anal Chem. 78:6717–6725.

68. Carey RJ, Damianopoulos EN, DePalma G (1994): Differential temporal dynamics of serum and brain cocaine: relationship to behavioral, neuroendocrine and neurochemical cocaine induced responses. Life Sci. 55:1711–1716.

69. Post RM (1980): Intermittent versus continuous stimulation: effect of time interval on the development of sensitization or tolerance. Life Sci. 26:1275–1282.

70. Downs AW, Eddy NB (1932): The effect of repeated doses of cocaine on the rat. The Journal of Pharmacology and Experimental Therapeutics. 46:199–200.

71. Post RM, Rose H (1976): Increasing effects of repetitive cocaine administration in the rat. Nature. 260:731–732.

72. Reith ME, Benuck M, Lajtha A (1987): Cocaine disposition in the brain after continuous or intermittent treatment and locomotor stimulation in mice. J Pharmacol Exp Ther. 243:281–287.

73. Stewart J, Badiani A (1993): Tolerance and sensitization to the behavioral effects of drugs. Behav Pharmacol. 4:289–312.

74. Rowland NE (2007): Food or fluid restriction in common laboratory animals: balancing welfare considerations with scientific inquiry. Comp Med. 57:149–160.

75. Samaha AN, Minogianis EA, Nachar W (2011): Cues paired with either rapid or slower self-administered cocaine injections acquire similar conditioned rewarding properties. PLoS One. 6:e26481.

76. Weeks JR (1962): Experimental morphine addiction: method for automatic intravenous injections in unrestrained rats. Science. 138:143–144.

77. Weeks JR, Davis JD (1964): Chronic Intravenous Cannulas for Rats. J Appl Physiol. 19:540–541.

78. Ricaurte GA, Seiden LS, Schuster CR (1984): Further evidence that amphetamines produce long-lasting dopamine neurochemical deficits by destroying dopamine nerve fibers. Brain Res. 303:359–364.

79. Ryan LJ, Linder JC, Martone ME, Groves PM (1990): Histological and ultrastructural evidence that D-amphetamine causes degeneration in neostriatum and frontal cortex of rats. Brain Res. 518:67–77.

80. Ryan LJ, Martone ME, Linder JC, Groves PM (1988): Continuous amphetamine administration induces tyrosine hydroxylase immunoreactive patches in the adult rat neostriatum. Brain Res Bull. 21:133–137.

81. Fortin GM, Bourque MJ, Mendez JA, Leo D, Nordenankar K, Birgner C, et al. (2012): Glutamate corelease promotes growth and survival of midbrain dopamine neurons. J Neurosci. 32:17477–17491.

82. Sanchez G, Varaschin RK, Bueler H, Marcogliese PC, Park DS, Trudeau LE (2014): Unaltered striatal dopamine release levels in young Parkin knockout, Pink1 knockout, DJ-1 knockout and LRRK2 R1441G transgenic mice. PLoS One. 9:e94826.

83. Varaschin RK, Osterstock G, Ducrot C, Leino S, Bourque MJ, Prado MAM, et al. (2018): Histamine H3 Receptors Decrease Dopamine Release in the Ventral Striatum by Reducing the Activity of Striatal Cholinergic Interneurons. Neuroscience. 376:188–203.

84. Paxinos G, Watson C (1986): The rat brain in stereotaxic coordinates. 2nd edn. Academic: New York, NY, USA.

85. Yorgason JT, Espana RA, Jones SR (2011): Demon voltammetry and analysis software: analysis of cocaine-induced alterations in dopamine signaling using multiple kinetic measures. J Neurosci Methods. 202:158–164.

